# Vitronectin defines a differentiation-restraining extracellular matrix state in myogenic cells

**DOI:** 10.64898/2026.01.14.699461

**Authors:** Tomoka Katayama, Anmi Matsumoto, Yoka Takazawa, Yuta Chigi, Daiji Okamura

**Affiliations:** Faculty of Agriculture, Kindai University; Independent researcher

**Keywords:** Vitronectin, Myogenic differentiation, Myoblast proliferation, Extracellular matrix, Serum-dependent regulation, Cell fate control, Replicative senescence, Integrin signaling, Skeletal muscle

## Abstract

Skeletal muscle differentiation is classically triggered by reducing serum-derived mitogenic cues; however, the extracellular matrix–associated factors that actively maintain a proliferative, differentiation-resistant state remain poorly defined. Here, we identify vitronectin (VN) as a key extracellular matrix gatekeeper that controls myogenic cell fate. VN is abundant in fetal bovine serum but largely absent from horse serum, rapidly declines upon differentiation induction, and selectively suppresses myogenic differentiation while sustaining proliferation through integrin αvβ3–dependent signaling. VN maintains growth factor receptor activity and cell-cycle progression even under differentiation conditions, thereby preventing myoblast commitment. These effects extend to three-dimensional culture and primary embryonic chicken myogenic cells, indicating evolutionary conservation. Importantly, VN progressively accumulates during long-term passaging, coinciding with replicative senescence and impaired differentiation, and inhibition of αvβ3 signaling partially restores myogenic competence in senescent cells. Together, our findings redefine a long-standing myoblast culture paradigm by identifying a serum-derived extracellular matrix protein as a decisive regulator linking extracellular environment, proliferative niche maintenance, and age-associated decline in myogenic competence.

## Introduction

The formation of skeletal muscle requires a finely regulated balance between myoblast proliferation and terminal differentiation. Although the transcriptional circuitry governing myogenesis has been extensively defined, extracellular cues play equally critical roles in determining whether myoblasts remain proliferative or commit to differentiation (1,2). Among these cues, the extracellular matrix (ECM) functions not only as a structural scaffold but also as a dynamic signaling platform that integrates adhesive interactions with growth factor–dependent pathways, thereby regulating cell survival, proliferation, and lineage progression (3,4).

Murine C2C12 myoblasts are widely used as an in vitro model of skeletal muscle differentiation (5). In this system, robust induction of myogenesis is classically achieved by switching cells from growth medium containing 10% fetal bovine serum (FBS) to differentiation conditions, most commonly based on horse serum (HS) (5,6). However, several studies have reported that efficient differentiation can also be induced under serum-free (7) or low-serum conditions using FBS alone, such as 0.5–1% FBS (6,8), without the addition of HS. These observations indicate that the transition to differentiation does not necessarily depend on a unique “differentiation-inducing factor” provided by HS; rather, a reduction in FBS itself can be sufficient to trigger myogenic differentiation (9,10). Historically, this transition has been largely attributed to the withdrawal of potent soluble mitogens present in FBS. In particular, fibroblast growth factor-2 (FGF2) has been widely recognized as a key regulator that sustains myoblast proliferation and suppresses the onset of differentiation, and its marked reduction in HS has been considered a major permissive event for myogenesis (1,11,12). By contrast, although IGF-1 plays important roles in myogenic precursor biology, their actions are highly context-dependent and neither has been consistently regarded as the primary determinant of this serum-dependent switch (13,14).

In parallel with soluble factors, serum composition also shapes the extracellular matrix environment encountered by myoblasts. Classical ECM components such as fibronectin, collagens, and laminins are well established as supportive substrates for myoblast adhesion, migration, fusion, and differentiation, and are generally permissive or even facilitative for myogenic commitment (15–19). These findings raise the possibility that, in addition to growth factors, serum-derived ECM-associated components may actively sustain a proliferation-competent and differentiation-resistant state. However, the identity of such ECM-linked suppressors of myogenic differentiation has remained unclear.

Another long-standing feature of the C2C12 system is its pronounced sensitivity to passage number (20). Extended culture is accompanied by replicative senescence, characterized by altered molecular profiles and a progressive decline in differentiation capacity (21,22). Although this phenomenon is widely recognized, the extracellular mechanisms linking prolonged culture, senescence-associated changes, and loss of myogenic competence remain poorly understood.

Here, we identify vitronectin (VN), a major serum-derived glycoprotein with extracellular matrix–associated adhesive functions (23,24), as a previously unrecognized regulator that connects these unresolved aspects of myoblast biology. We show that VN uniquely suppresses both morphological and molecular hallmarks of myogenic differentiation while sustaining proliferation, growth factor receptor signaling, cell survival, and cell-cycle progression through integrin-dependent mechanisms. VN is abundant in FBS but largely absent from HS, and its selective depletion from serum markedly enhances myogenic differentiation. In parallel, under serum-free conditions, the presence of VN is sufficient to maintain myoblast proliferation and an undifferentiated state. Together, these complementary loss- and gain-of-function observations indicate that the reduction of serum-derived VN constitutes a decisive event that permits myogenic commitment. Moreover, VN progressively accumulates during long-term culture, coinciding with replicative senescence and impaired myogenic differentiation. Importantly, pharmacological inhibition of integrin αvβ3 signaling partially restores differentiation in senescent myoblasts, indicating that VN–integrin signaling is not only correlative but also functionally contributes to senescence-associated loss of myogenic competence. Together, these findings position vitronectin as an extracellular gatekeeper of myogenic cell fate and suggest that its accumulation contributes to senescence-associated loss of myogenic plasticity.

## Materials and Methods

### Culture of C2C12 cells for proliferation and differentiation

C2C12 cells (an immortalized mouse myoblast cell line) (5,6) were cultured in growth medium (GM) consisting of DMEM (FUJIFILM Wako, 043-30085) supplemented with 10% fetal bovine serum (FBS; Gibco, 10270-106) and 1x penicillin-streptomycin mixed solution (P/S; FUJIFILM Wako, 168-23191). Cells were passaged every 3-4 days using TrypLE (Gibco, 12604-013) at a split ratio of 1:5. C2C12 myoblasts are capable of differentiating into multinucleated myotubes under low serum condition (6). For induction of differentiation, 3.0 x 10^4^ cells were seeded into each well of a 4-well plate (SPL Life Sciences, SPL-30004) in GM. After 2-3 days, when cells reached 90-100% confluence, the medium was replaced with differentiation medium (DM) consisting of DMEM supplemented with 2% horse serum (HS; Sigma, H1138) and 1x P/S. To extend the differentiation culture for 5 days, the DM was refreshed on day 3.

Cyclic RGDfV (cRGDfV) is an RGD-containing peptide antagonist with specificity for αvβ3 integrin and is commonly used to inhibit the binding of cells to RGD-dependent ECM proteins such as vitronectin (SCP0111, Sigma-Aldrich). In this study, cRGDfV was dissolved in dimethyl sulfoxide (DMSO) and used at a final concentration of 1 µM, while DMSO was added at 0.04% as a vehicle control.

### Pre-coating with extracellular matrix

Prior to seeding C2C12 cells, the extracellular matrix (ECM) was coated onto individual wells of 4-well culture plate at the desired coating density, followed by incubation at 37 C° for 2 h and aspiration of excess solution before use. The following ECM coating density were used: fibronectin, 2.35 μg/cm^2^ (FUJIFILM, 063-05591); vitronectin, 0.17∼2.8 μg/cm^2^ (FUJIFILM, 220-02041); laminin 511-E8 fragment iMatrix-511, 0.5 μg/cm^2^ (Nippi, 892011); Matrigel, 2% (Corning, 354234); Gelatin, 0.2% gelatin (Sigma, G1890); and collagen type I, 0.2% (Nippi, ASC-1-100). Following the optimization as shown in Supplementary Figure 2 and 4, the coating density of vitronectin was determined as 0.7 μg/cm^2^ for a serum-containing medium and 1.4 μg/cm^2^ for a serum-free medium.

### Immunofluorescence

Cells cultured in 4-well plates were fixed with freshly prepared 4% paraformaldehyde in Dulbecco’s phosphate-buffered saline (PBS; FUJIFILM Wak, 041-20211) for 15 min at room temperature, followed by permeabilization and blocking in PBS containing 1% Triton X-100 (FUJIFILM Wako, A16046), 1% BSA (Sigma, A3059), and 10% newborn calf serum (CS) (biowest, S0750) for 1 h at room temperature. Cells were then incubated overnight at 4℃ with primary antibodies diluted in PBS containing 1% BSA, 1% CS, and 0.1% Triton X-100. Following three washes with PBS containing 0.1% Triton X-100 (PBST), cells were incubated for 2 h at room temperature or overnight at 4 °C with fluorophore-conjugated secondary antibodies diluted 1:500 in PBST containing 1% BSA and 1% CS. Nuclei were counterstained with DAPI (Sigma-Aldrich, D9542). Following three washes with PBST, samples were mounted in PBS for imaging. Fluorescence images were obtained using a BZ-X710 microscope (Keyence).

### Antibodies for immunofluorescence

As a primary antibody, Anti-Myosin Heavy Chain (MHC) antibody (MF20, 1:500; R&D Systems, MAB4470), Anti-Myogenin antibody (1:500; Abcam, ab124800), Anti-p-Histone H3 antibody (C-2, 1:1000; Santa Cruz, sc-374669), Anti-Cleaved Caspase-3 antibody (Asp175) (5A1E, 1:500; Cell Signaling Technology, #9664), Anti-Ki-67 (1:500; Abcam, ab16667). As a secondary antibody, Goat anti-mouse IgG (H+L), Alexa Fluor 488 (Abcam, ab150113); Goat anti-mouse IgG (H+L), Alexa Fluor 594 (Abcam, ab150116); Goat anti-rabbit IgG (H+L), Alexa Fluor 594 (Abcam, ab150080).

### Immunoblot analysis

Cells were lysed in SDS sample buffer containing 1M Tris-HCl (pH 6.8; FUJIFILM Wako, 015-20093), glycine (FUJIFILM Wako, 077-00735), 10% SDS (FUJIFILM Wako, 192-13981), 10 mg/mL bromophenol blue (BPB) (nacalai tesque, 05808-61), and 2-Mercaptoethanol (nacalai tesque, 21438-82). Protein concentration was determined using the Qubit Protein Assay Kit (Invitrogen, Q33211). Proteins were separated by SDS-PAGE using a 12% separating gel and a 4.5% stacking gel prepared from 30% (w/v) acrylamide/bis-acrylamide solution (29:1; FUJIFILM Wako, 015-25635), 1 M Tris-HCl (pH 8.8 for the separating gel, pH 6.8 for the stacking gel), 10% SDS, 10% (w/v) ammonium persulfate (APS; nacalai tesque, 02634-34), and TEMED (nacalai tesque, 33401-72). Electrophoresis was performed in 1× SDS running buffer, and proteins were transferred onto a PVDF membrane (MERCK, IPVH0010) using the HorizeBLOT 2M system (ATTO, WSE-4025) at 18 V/153 mA for 30 min. Membranes were blocked for 1h at room temperature with either 0.3% skim milk in TBST (TBS + 0.1% Tween-20 (FUJIFILM Wako, 166-21213) or EzBlock Chemi (ATTO, AE-1475). After blocking, membranes were incubated overnight at 4 °C with primary antibodies diluted in Can Get Signal^®^ Immunoreaction Enhancer Solution 1 (TOYOBO, NKB-101). Following three washes with TBST, membranes were incubated with HRP-conjugated secondary antibodies (1:10000 dilution) in Can Get Signal^®^ Immunoreaction Enhancer Solution 2 (TOYOBO, NKB-101) for 3 h at room temperature or overnight at 4 °C. Following three washes with TBST, signal detection was performed using ECL Start (Cytiva, RPN3243), ECL Prime (Cytiva, RPN2232), or ECL Select (Cytiva, RPN2235) and imaged with a LuminoGraph I imaging system (ATTO, WSE-6100). Molecular weights were verified using the EzProtein Ladder (ATTO, WSE-7020). To compare vitronectin abundance among different serum sources, FBS, adult bovine serum (ABS)(B9433, Sigma-Aldrich), and HS were analyzed by immunoblotting. Each serum sample was normalized by total protein content, and 1 µg of total protein from each serum was applied.

### Antibodies for immunoblot analysis

As a first antibody, Anti-MyoD (5.8A, 1:5000; Novus Biologicals, NB100-56511), anti-Myogenin (1:10000; Abcam, ab124800), Anti-MHC (MF20, 1:5000; R&D Systems, MAB4470), Anti-β-Actin (C4, 1:5000; Santa Cruz, sc-47778), Anti-Stat3 (D3Z2G, 1:5000; Cell Signaling Technology, #12640), Anti-phospho-Stat3 (Tyr705, D3A7, 1:5000; Cell Signaling Technology, #9145), Anti-ERK1/2 (C-9, 1:5000; Santa Cruz, sc-514302), Anti-phospho-ERK (E-4, 1:5000; Santa Cruz, sc-7383), Anti-p38α/β MAPK (A-12, 1:5000; Santa Cruz, sc-7972), anti-phospho-p38 (E-1, 1:5000; Santa Cruz, sc-166182), Anti-p21 Waf/Cip1 (F-5, 1:5000; Santa Cruz, sc-6246), Anti-p27 Kip2 (F-8, 1:5000; Santa Cruz, sc-1641), Anti-FGF Receptor 1 (D8E4, 1:5000; Cell Signaling Technology, #9740), Anti-p-FGF Receptor 1 (Try653/654) (D4X3D, 1:5000; Cell Signaling Technology, #52928), Anti-Insulin Receptor β (4B8, 1:5000; Cell Signaling Technology, #3025), Anti-p-IGF-1 Receptor β (Tyr1135/1136)/Insulin Receptor β (Try1150/1151) (19H7, 1:5000; Cell Signaling Technology, #3024), Anti-Vitronectin 65/75 (D-8, 1:5000; Santa Cruz, sc-74484), Anti-p16^INK4A^ (F-12, 1:5000; Santa Cruz, sc-1661), p-χHistone H2A.X (Ser 139, 1:5000; Santa Cruz, sc-517348). As a secondary antibody, Anti-mouse IgG (H+L chain), pAb-HRP (MBL, 330); Anti-rabbit IgG (H+L chain), pAb-HRP (MBL, 458).

### RNA preparation and real-time PCR

Total RNA was extracted using RNeasy Mini Kit (QIAGEN, 74104) according to the manufacturer’s instructions. cDNA was synthesized using the ReverTra Ace qPCR RT Master Mix (TOYOBO, FSQ-201), and real-time PCR was performed with the THUNDERBIRD SYBR qPCR Mix (TOYOBO, QPS-201) on a Mic qPCR Cycler (bio molecular systems). Gene expression levels were normalized to mouse *Hprt* expression and calculated using the comparative CT (ΔΔCt) method. Primer sequences were as follows:

*Hprt*-F (CAGTCCCAGCGTCGTGATTA), *Hprt*-R (AGCAAGTCTTTCAGTCCTGTC), *Vitronectin*-F (CCCCTGAGGCCCTTTTTCATA), *Vitronectin*-R (CAAAGCTCGTCACACTGACA).

### Immunodepletion of vitronectin from FBS

rProtein A Sepharose^TM^ Fast Flow beads (Cytiva, 17127901) were gently rotated for 2 h at room temperature to ensure homogeneous suspension. A 100 μL aliquot of beads was washed once with KMH buffer (100 mM KCl [FUJIFILM Wako, 169-03542], 2.5 mM MgCl_2_ [FUJIFILM Wako, 135-00165], and 20 mM HEPES-KOH [pH 7.5; nacalai tesque, 15639-84]) by centrifugation (300 x g, 5 min). Beads were then incubated with 10 μL of anti-vitronectin 65/75 antibody (D-8; Santa Cruz, sc-74484) and control antibody (Santa Cruz, sc-3877) overnight at 4℃. Following two washes with KMH buffer (300 x g, 5 min), 200 μL of FBS (Gibco, 10270-106) was added to the antibody-bound beads, and the mixture was rotated overnight at 4℃. The supernatant was collected and filtered through 0.2 µm membrane filter (Advantec, 13CP020AS) to obtain vitronectin-depleted FBS and control FBS.

### Culture in serum-free medium

The detailed composition of the serum-free media used in this study, including DA-X, is described below. The DA-X medium (25), a fully serum-free medium used in this study, was based on DMEM (high glucose) with L-glutamine, phenol red, and sodium pyruvate (FUJIFILM Wako, 043-30085), supplemented with Insulin–Transferrin–Selenium–Ethanolamine (ITS-X) (1×; Gibco, 51500056), bovine serum albumin (BSA) (5 mg/mL; Sigma, A3059), L-glutamine (2 mM; FUJIFILM Wako, 073-05391), and penicillin–streptomycin mixed solution (1×; FUJIFILM Wako, 168-23191). Where indicated, leukemia inhibitory factor (LIF) was supplied at 0.1% (v/v) using conditioned medium derived from COS cells transiently transfected with a LIF expression vector (see below). In addition, water-soluble cholesterol was added to the medium at a final concentration of 10 µM, and β-mercaptoethanol (2-ME) (Nacalai Tesque, 21438-82) was added at a final concentration of 100 µM. All medium components were pre-mixed and stored at 4 °C. However, the water-soluble cholesterol solution was added freshly immediately before use to prevent crystallization in the pre-mixed medium m (26). The preparation of water-soluble cholesterol is described below. For selected experiments, vitronectin was coated at a density of 1.4 µg/cm², and recombinant human IGF-1 (2 ng/mL; PeproTech, #100-11) and FGF-2 (100 ng/mL; PeproTech, #100-18B) were added as indicated.

### Preparation of LIF-conditioned medium

LIF-conditioned medium was prepared from COS cells transiently transfected with the pCAGGS–LIF expression vector (27). Briefly, COS cells were seeded in 10-cm dishes and cultured to approximately 70–80% confluence, followed by transfection using Lipofectamine 3000 (Thermo Fisher Scientific) according to the manufacturer’s instructions. On the following day, transfected cells were detached with trypsin and passaged at approximately 1:4 dilution into four new 10 cm dishes. Culture supernatants were collected on days 3 and 4, filtered through a 0.22 µm membrane filter, and stored at −20 °C until use. The conditioned medium was added to serum-free cultures at a 1:1000 dilution as a source of LIF. As a control, an equivalent volume of conditioned medium collected from COS cells transfected with an empty vector was added in parallel.

### Preparation of water-soluble cholesterol solution

For the preparation of 50 mM cholesterol/EtOH stock solution, 19.3 mg of cholesterol (Sigma, C8667) was dissolved in 1 mL of 100% EtOH by incubation at 80°C. For water-solubilization of the cholesterol/EtOH solution, 42 mg of Methyl-β-cyclodextrin (Sigma-Aldrich, 332615) was added to the 1 mL of 5 mM cholesterol/EtOH solution. The solution was stored at -30°C before use.

### Rotary suspension spheroid culture

Cells were seeded at a density of 6.0 x 10^3^ cells per well in 96-well U-bottom plate (Thermo Fisher Scientific, 174925) and incubated for one day in 100 µL of GM. On the following day, formed spheroids were carefully transferred using a MICROCAPS pipette (Drummond, 1-000-0500) under a stereomicroscope to a 4-well plate (SPL Life Sciences, SPL-30004) with 500 µL of DM. The plates were rotated at 150 rpm on a rotary shaker (Waken B TECH, WB-T101SRC) as a rotary suspension spheroid culture. After completing the immunofluorescence staining, the spheroids were transferred to APS-coated glass slides (MATSUNAMI, SAPS-01) and mounted in VECTASHILDE^R○^ mounting medium (Vector, H-1200), after which the excess PBST was removed. The slides were then covered with NEO cover glass (MATUNAMI, C024501) and sealed with a nail topcoat. Images were acquired using a fluorescence microscope (Keyence, BZ-X710) and a confocal laser scanning microscope (Olympus, FV-3000).

### Primary culture of embryonic chick myogenic cells

As described previously (28), thigh muscle tissue was dissected using fine forceps from 10-day-old chick embryos obtained from a certified poultry supplier. The femurs were surgically removed, and the isolated muscle tissue was finely minced and washed twice with PBS. The minced tissue was then digested in HBSS (nacalai tesque, 09735-75) containing 2 mg/mL collagenase type II (Worthington Biochemical Corporation, WOR-CLS2) at 37°C for 40–60 min, with gentle pipetting every 10–15 min to facilitate cell dissociation. The resulting cell suspension were filtered through a 40 µm cell strainer (Corning, 431750) to obtain single cells. Approximately 5.0 × 10⁵ cells were seeded per well in 4-well plates pre-coated with either 0.2% gelatin (control) or 0.7 μg/cm² vitronectin (VN) in GM to ensure stable attachment. On the following day, the medium was replaced with DM, and cells were induced to differentiate for approximately 6 h. Cultured embryonic chick myogenic cells were then subjected to subsequent analyses. All animal experiments were conducted in accordance with the ethical guidelines of Kindai University and were reviewed and approved by the Kindai University Animal Care and Use Committee.

### Statistical analysis

All statistical analysis was performed using one-way ANOVA. When overall significance was observed, post hoc comparisons between groups were evaluated using Student’s t-test. p values <0.05 were considered statistically significant.

## Results

### Vitronectin suppresses myogenic differentiation through integrin signaling

C2C12 myoblasts proliferate stably and robustly in growth medium (GM; DMEM supplemented with 10% FBS), whereas switching to differentiation medium (DM; DMEM supplemented with 2% HS) rapidly induces myogenic differentiation accompanied by cell fusion and robust induction of myotube-specific marker genes (Figure 1a). Among the major extracellular matrix (ECM) components known to be present in FBS, fibronectin, collagen, and laminin have been extensively characterized and are generally recognized to support myoblast adhesion, proliferation, and differentiation. In contrast, the functional contribution of vitronectin (VN) to myogenic regulation has remained poorly defined.

**Figure 1.**
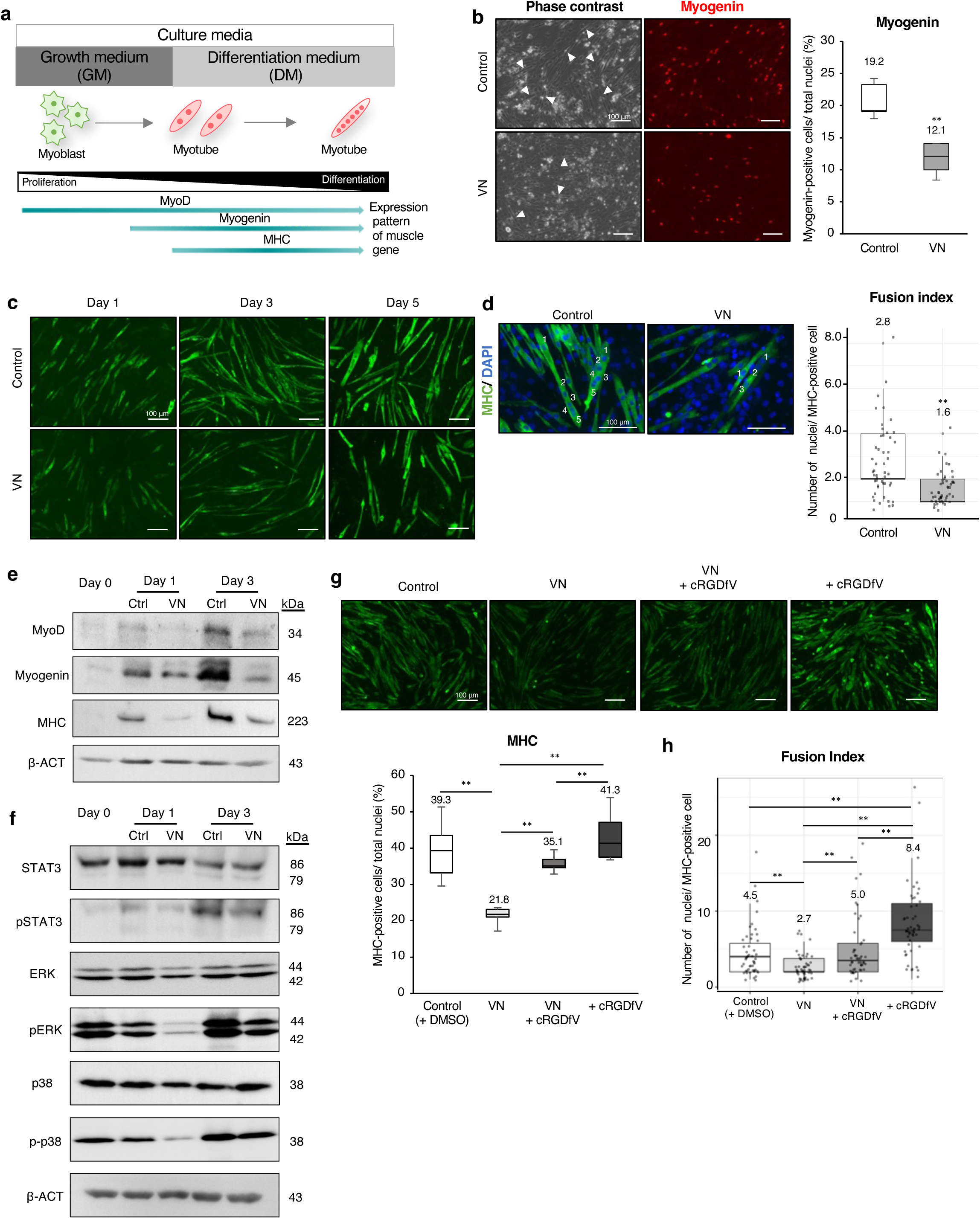
Inhibitory effects of vitronectin on the myogenic differentiation in mouse C2C12 myoblasts. **a,** Schematic representation of the myogenic differentiation processes in C2C12 myoblasts dependent on culture medium and expression dynamics of representative muscle differentiation markers. Proliferation of myoblasts requires growth medium (GM; 10% fetal bovine serum (FBS) supplemented medium), whereas differentiation medium (DM; 2% horse serum (HS) supplemented medium) induces rapid differentiation into multinucleated myotubes and loss of proliferative capacity. **b,** Representative phase contrast and immunofluorescence images showing expression of the differentiation marker Myogenin on day 3 in DM. White arrowheads indicate representative differentiating myotubes. Differentiation index quantified by the proportion of myogenin-positive cells relative to total nuclei is shown in the graph. (Control: n = 1652, Vitronectin: n = 1746) **c,** Time-course observation of myotube differentiation by immunofluorescence for myosin heavy chain (MHC). **d,** Fusion index quantified by the number of nuclei per MHC-positive myotube on day 3 in DM shown in the graph (Control: n=50, Vitronectin: n=50 in MHC-positive cells). The numbers shown in the immunofluorescence images indicate of nuclei in the representative MHC-positive myotube. **e,** Immunoblot analysis of C2C12 cells for the representative myogenic differentiation markers: MyoD, Myogenin, and MHC. **f,** Immunoblot analysis of C2C12 cells for non- and phosphorylated form of STAT3, ERK1/2, and p38 MAPK. **g-h**, Immunofluorescence analysis of MHC expression after 3 days in DM under the indicated conditions: control, VN, VN plus cRGDfV, and cRGDfV alone. Quantification shows the proportion of MHC-positive cells relative to total nucle as differentiation indexi (g) and fusion index (h). Scale bar: 100 µm. Nuclei were counterstained with DAPI. Data in the graphs are presented as mean ± s.d. Statistical significance determined using Student’s t-test; \*\**p < 0.01*.

C2C12 myoblasts proliferate in growth medium (GM; DMEM supplemented with 10% FBS) but undergo rapid myogenic differentiation upon transfer to differentiation medium (DM; DMEM supplemented with 2% HS) (Figure 1a). While several ECM components present in FBS, including fibronectin, collagen, and laminin, are known to support myogenic differentiation, the role of vitronectin (VN) in myoblast fate regulation has remained unclear.

To address this, we compared the effects of VN with those of representative ECM substrates. Under differentiation conditions, C2C12 cells cultured on fibronectin, laminin, collagen type I, or other commonly used adhesive matrices formed multinucleated myotubes and robustly expressed myogenic markers. In contrast, cells cultured on VN-coated surfaces exhibited significantly reduced myotube formation, with fewer myogenin- and MHC-positive cells and a low fusion index, remaining predominantly mononuclear (Figures 1b–d; Figures S1, S2).

Immunoblot analysis further revealed that VN markedly attenuated expression of MyoD, myogenin, and MHC, accompanied by reduced phosphorylation of STAT3 (29–31), ERK1/2 (32), and p38 MAPK (33,34) signaling pathways implicated in myogenic differentiation (Figures 1e, f). These results identify VN as a unique ECM component that suppresses both morphological and signaling programs required for myogenic differentiation.

Because VN primarily signals through integrin αvβ3 (35)., we next examined whether this pathway mediates the inhibitory effects of VN Pharmacological inhibition of αvβ3 using the competitive peptide cRGDfV (35,36) significantly restored myotube formation and MHC expression in VN-coated cultures (Figure 1g). Notably, cRGDfV treatment also enhanced differentiation under control conditions, suggesting that basal αvβ3 signaling restrains myogenic differentiation. Quantification of fusion index confirmed a robust rescue of differentiation upon αvβ3 inhibition (Figure 1h).

Together, these findings demonstrate that VN suppresses myogenic differentiation via integrin αvβ3-dependent signaling, establishing VN as a differentiation-restraining ECM component in myoblasts.

### Vitronectin sustains growth factor signaling and proliferation under differentiation conditions

Given that VN suppresses myogenic differentiation, we next asked whether VN preserves proliferative signaling programs that are normally attenuated upon differentiation induction. This question is grounded in the developmental principle that entry into terminal differentiation is typically coupled to cell-cycle exit and loss of mitogenic signaling (37–39).

To directly assess proliferative responses under differentiation conditions, C2C12 myoblasts were seeded at low density and cultured on VN-coated surfaces (Figure 2a). Cell counts revealed a clear, VN-dependent increase in total cell number compared with control cultures (Figure 2b; Figures S1b, S2c). Consistently, immunofluorescence analysis showed an increased proportion of phosphorylated histone H3–positive cells on VN, indicating sustained mitotic activity despite differentiation cues (Figure 2c). In parallel, the frequency of cleaved caspase-3–positive cell was reduced, suggesting enhanced cell survival under VN-coated conditions (Figure 2d).

**Figure 2.**
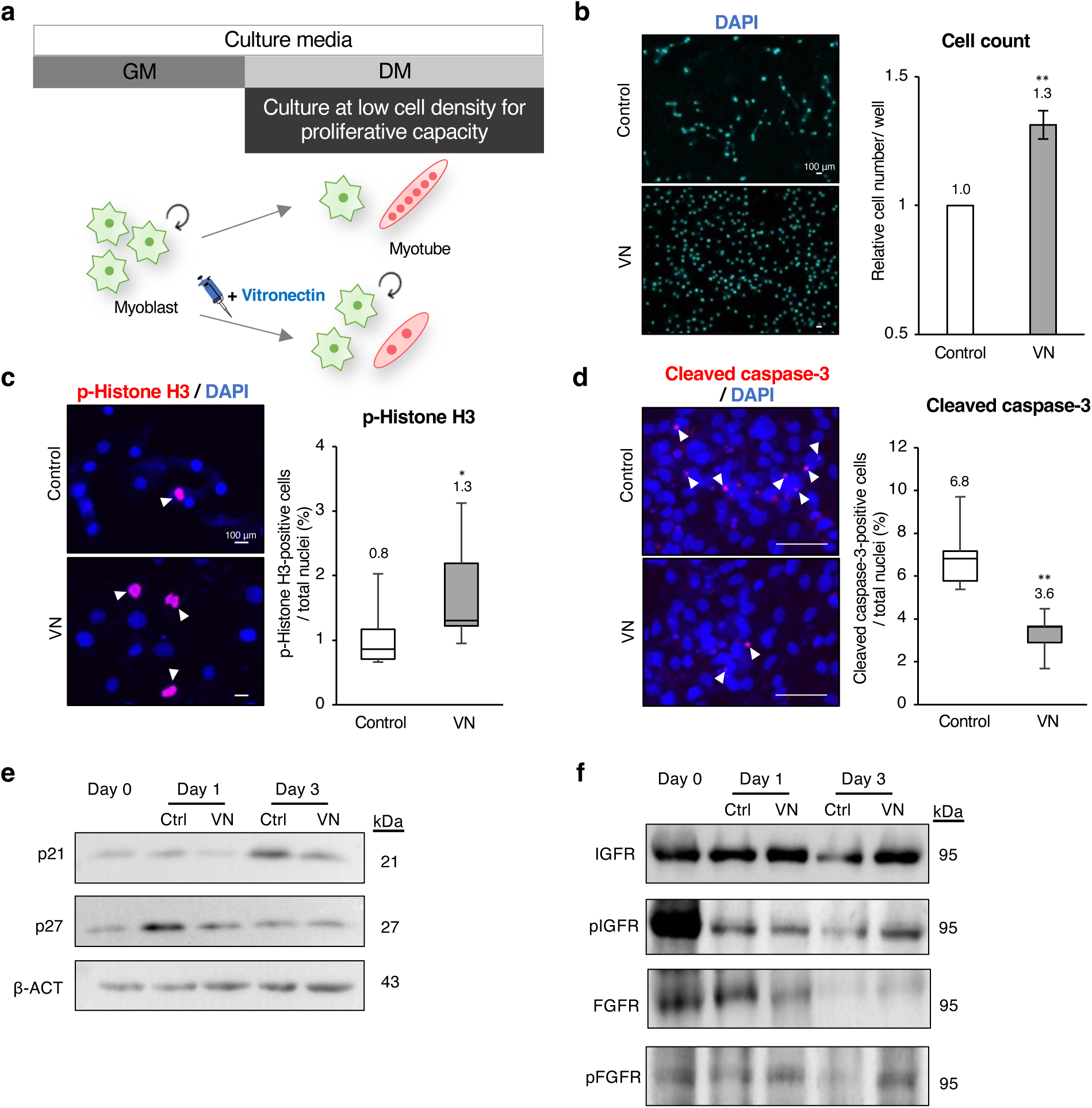
Vitronectin ensures the proliferation of C2C12 myoblasts under DM conditions. **a,** Schematic representation of the experimental workflow to evaluate the proliferative capacity of C2C12 myoblasts cultured at low cell density in supplementation with vitronectin (GM: growth medium, DM: differentiation medium). **b,** Representative nuclei images stained with DAPI and the relative cell number of C2C12 cells counted on day 3 of differentiation shown in the graph (n = 6). **c,** Representative immunofluorescence images for phospho-Histone H3 (pH3) on day 3 of differentiation and the proportion of pH3-positive cells relative to total nuclei shown in the graph (Control: n = 1345, Vitronectin: n = 2632). **d,** Representative immunofluorescence images for cleaved caspase-3 on day 2 of differentiation. The ratio of apoptotic cells relative to total nuclei is shown in the graph (Control: n = 2499, Vitronectin: n = 2345). **e,** Western blot analysis of C2C12 cells for p21 and p27, cell cycle inhibitors. **f,** Immunoblot analysis of C2C12 cells for non- and phosphorylated insulin-like growth factor receptor (IGFR) and fibroblast growth factor receptor (FGFR), key growth factor receptors involved in the proliferation of C2C12 cells. Scale bar: 100 µm. White arrowheads indicate representative positive cells for p-Histone H3 (c) and cleaved caspace-3 (d). Data are presented as mean ± s.d. Statistical significance determined using Student’s t-test; \*\**p < 0.01*, \**p < 0.05*.

To further explore how VN promotes proliferation, we examined whether it interferes with cell cycle exit, which is a prerequisite for myogenic differentiation. Immunoblot analysis showed that the cell cycle inhibitors p21 (40,41) and p27 (42) were downregulated under VN-coated conditions (Figure 2e). This finding suggests that VN suppresses cell cycle arrest, thereby maintaining the proliferative capacity of myoblasts even under differentiation conditions.

The myoblasts continued to proliferate; thus, we investigated whether VN maintains activation of growth factor signaling pathways that support myoblast proliferation. Insulin-like growth factor receptor (IGFR) signaling stimulates the PI3K/Akt cascade to promote myoblast proliferation (43,44), whereas fibroblast growth factor receptor (FGFR) signaling suppresses differentiation while enhancing proliferation (45,46). Both IGFR and FGFR exhibited increased phosphorylation under VN-coated conditions, indicating activation of these growth factor pathways (Figure 2f).

Together, these results demonstrate that VN sustains growth factor receptor signaling and proliferative capacity under differentiation conditions, thereby maintaining myoblasts in a growth-competent state that resists myogenic commitment.

### Vitronectin suppresses myogenic differentiation and sustains proliferation in a three-dimensional culture context

Because myogenic differentiation and proliferation are strongly influenced by cell–cell interactions and tissue architecture, we next asked whether the differentiation-suppressive and proliferation-supportive effects of VN observed in two-dimensional cultures are preserved in a three-dimensional context. To address this, we employed a suspension-based spheroid culture system that allows myoblasts to self-organize into compact three-dimensional aggregates (Figure 3a).

**Figure 3.**
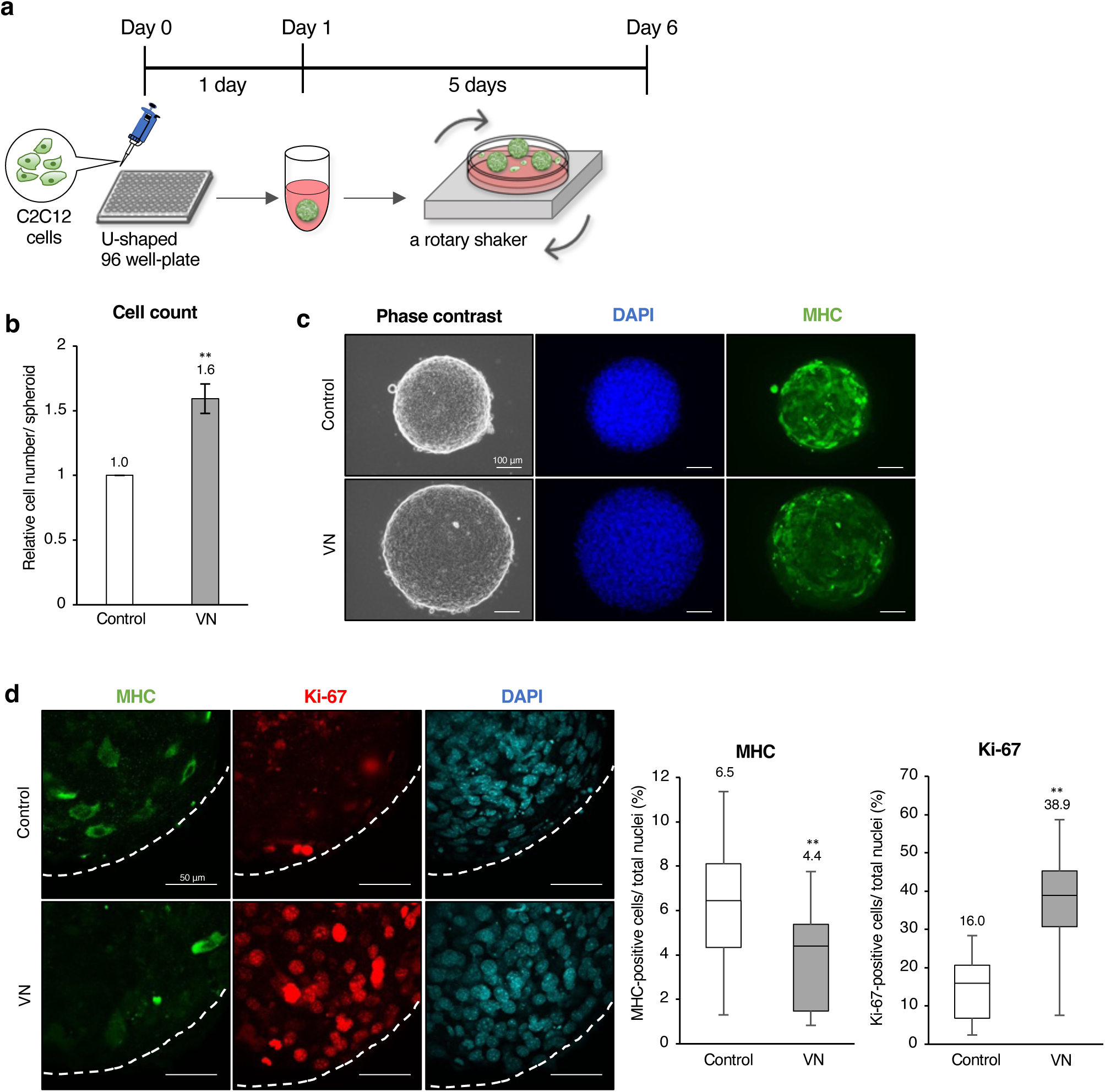
Vitronectin enhances myoblast proliferation and suppresses differentiation in rotary suspension spheroid culture. **a,** Schematic representation of the rotary suspension spheroid culture system. C2C12 cells were first aggregated under static conditions in a low-attachment U-shaped 96-well plate for 1 day, then transferred to suspension culture on an orbital shaker inside an incubator (37 °C, 150 rpm.) for another 5 days. **b,** Quantification of total cell numbers per spheroid on day 6 (n = 5). **c,** Representative phase-contrast and immunofluorescence images of spheroid for MHC and DAPI. Scale bar: 100 µm. **d,** Representative confocal images of spheroid stained for MHC, Ki-67, and DAPI on day 6. Quantification of the proportion of MHC- or Ki-67-positive cells relative to total nuclei is shown in the graphs (Control: n = 1537, Vitronectin: n = 1522). Scale bar: 50 µm. Data are presented as mean ± s.d. Statistical significance was determined using Student’s t-test; \*\**p < 0.01*.

Under VN-supplemented media conditions, myoblast spheroids exhibited a significant increase in overall cell number (Figure 3b) and spheroid size compared with the control (Figure 3c). Images of phase contrast and DAPI staining confirmed the enlarged aggregates, while immunofluorescence staining revealed reduced expression of the myotube marker, MHC (Figure 3c). Confocal imaging further demonstrated that MHC expression was diminished, whereas the proliferation marker Ki-67-positive cells were remarkably increased in VN-treated spheroids relative to the control (Figure 3d). These findings indicate that VN suppresses myogenic differentiation and maintains a proliferation-competent state not only on planar substrates but also within a three-dimensional cellular architecture that more closely resembles tissue-like organization.

### Serum-dependent vitronectin abundance governs the balance between myoblast self-renewal and differentiation

Since our results demonstrated that VN suppresses myogenic differentiation and promotes proliferation in both 2D and 3D culture contexts (Figures 1-3), we next examined whether serum-dependent differences in VN abundance regulate the balance between myoblast self-renewal and differentiation. Because serum replacement from FBS to HS is a critical step for initiating C2C12 differentiation, we hypothesized that reduced VN levels in HS may act as a physiological cue for the transition from proliferation to differentiation. To test this, we compared VN protein levels among FBS, adult bovine serum (ABS), and HS by immunoblot analysis (Figure 4a, Figure S3). VN was highly abundant in FBS but nearly undetectable in HS, and this difference was not species-dependent, as VN levels were also considerably lower in ABS (Figure 4a).

**Figure 4.**
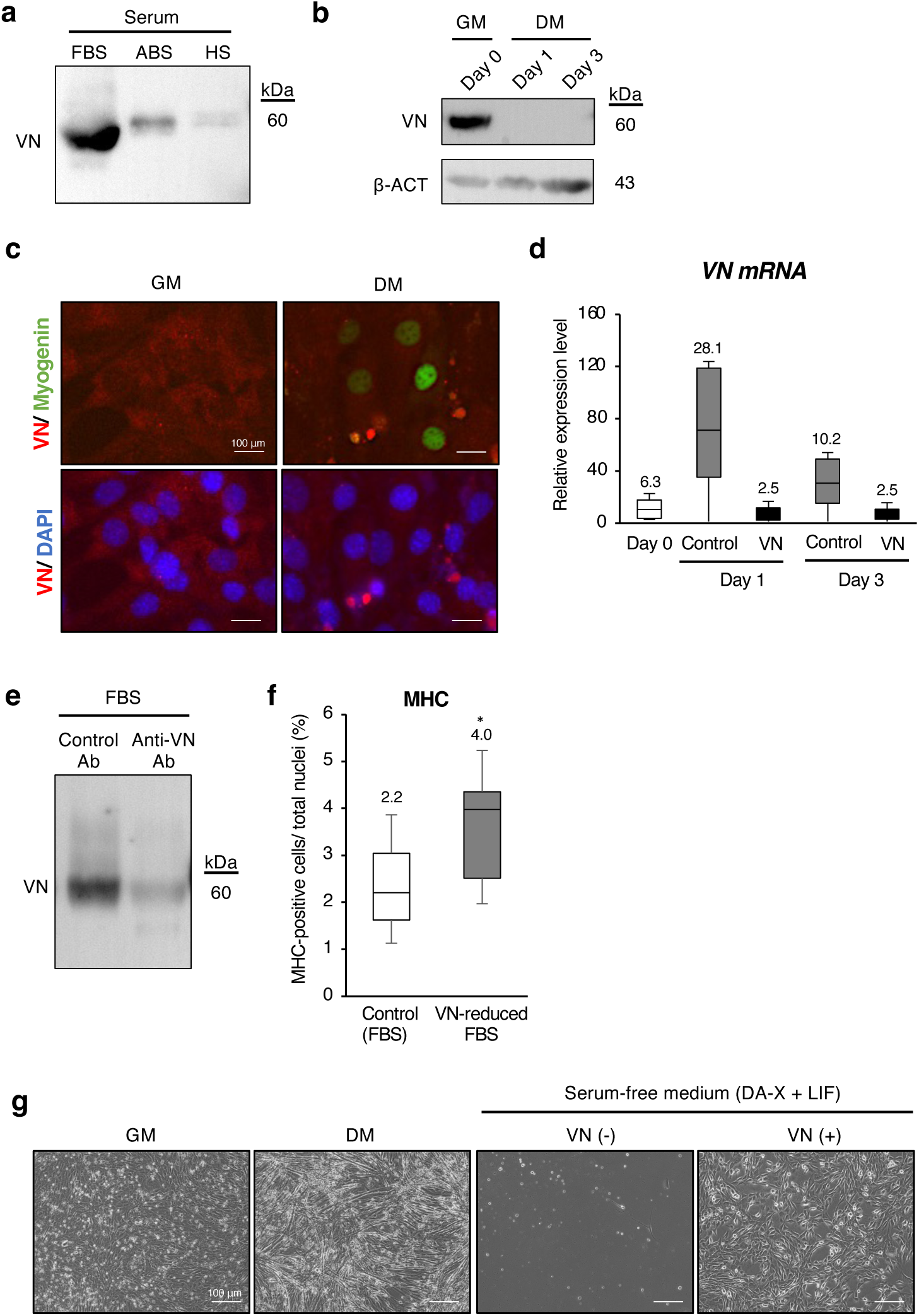
Vitronectin abundance underlies the inhibitory effects of FBS on myoblast differentiation. **a,** Immunoblot analysis of VN protein levels in fetal bovine serum, adult bovine serum, and adult horse serum. **b,** Immunoblot analysis of VN protein on day 0 (d0; before differentiation induction), day 1, and day 3 (d1 and d3; 1 and 3 day after differentiation induction). **c,** Representative immunofluorescence images showing the level and localization of VN protein in growth medium (GM) and day 1 in differentiation medium (DM). **d,** qPCR analysis for transcription of VN gene on d0, d1, and d3 after differentiation induction (n = 3). **e,** Immunoblot analysis for confirmation of immunodepleting of vitronectin from FBS. Vitronectin was depleted from FBS using control and anti-vitronectin antibody immobilized on Protein A Sepharose. **f,** Quantification of the inhibitory effect on myoblast differentiation with control and VN-immunodepleted FBS by immunofluorescence for MHC. (Control FBS: n = 1688, VN-reduced FBS: n = 1663 in counted nuclei number). **g,** Representative phase-contrast images of C2C12 cells cultured in GM, DM, and a serum-free DA-X medium supplemented with LIF on plates coated with or without VN coating. Images were captured on day 5 in each condition. Scale bar: 100 µm. Data are presented as mean ± s.d. Statistical significance was determined using one-way ANOVA in **d** and Student’s t-test in **f**; \**p < 0.05*.

We then examined VN dynamics during C2C12 differentiation. VN was highly detectable at day 0 in GM but its levels decreased sharply by day 1 in DM (Figure 4b). Immunofluorescence staining confirmed a corresponding reduction of VN protein of the cells in differentiation conditions (Figure 4c). qPCR analysis further demonstrated that the VN protein detected at day 0 was not attributable to endogenous transcription, as *VN* mRNA remained low compared with differentiating controls (Figure 4d). These findings suggest a feedback mechanism in which extracellular, FBS-derived VN suppresses cellular VN production, indicating that myogenic regulation under standard culture conditions is primarily governed by exogenous VN rather than cell-derived VN (Figures 4a–d).

Consistent with this, selective immunodepletion of VN from FBS significantly enhanced myotube formation even under 5% FBS conditions, which are normally sufficient to maintain myoblasts in an undifferentiated state (Figures 4e, 4f), demonstrating that reducing serum-derived VN is a decisive event that promotes differentiation.

### Vitronectin establishes a canonical growth factor–independent proliferative niche under serum-free conditions

We next examined whether vitronectin could sustain myoblast proliferation under serum-free conditions independently of canonical soluble growth factors. Historically, FGF2 and IGF-1 (47,48) have been regarded as critical serum-derived factors that maintain myoblast proliferation and suppress differentiation. To directly test whether these signals are dispensable, we assessed the effect of VN in defined serum-free media.

Building on previously established serum-free DA-X medium (25)(26), originally optimized to suppress differentiation and sustain proliferation in multiple cancer cell lines and mouse embryonic stem cells, respectively, we optimized these basal conditions to evaluate the contribution of VN to myoblast fate control (Figure S4). Whereas C2C12 cells typically cease proliferation and undergo differentiation or apoptosis upon serum withdrawal (Figure 4g), cells cultured on VN-coated surfaces in serum-free medium supplemented with leukemia inhibitory factor (LIF) remained viable and proliferative, while resisting spontaneous differentiation, even in the absence of exogenously supplied FGF2 or IGF-1 (Figure 4g). Consistent with these observations, systematic optimization of defined serum-free conditions revealed that vitronectin provides most efficient extracellular scaffold for myoblast survival and proliferation, with LIF being essential for long-term expansion, whereas additional components such as cholesterol and redox modulation exerted supportive but non-essential effects (Figure S4).

These findings demonstrate that VN, in combination with LIF, establishes an extracellular matrix–based proliferative state that is fundamentally independent of canonical soluble mitogens, thereby redefining the long-standing growth factor–centric model of myoblast maintenance.

### Vitronectin suppresses myogenic differentiation and sustains proliferation in primary chicken embryonic myoblasts

To determine whether the functions of VN observed in murine C2C12 cells are conserved across species, we examined its effects in primary myogenic cells isolated from 10-day-old chicken embryos. Because embryonic myoblast preparations often contain pre-fused myotubes and undergo rapid differentiation, we employed a short-term differentiation assay (28).

Under these conditions, VN supplementation significantly reduced the fusion index after 6 hours of differentiation, indicating suppressed myotube formation in primary chicken myoblasts (Figure 5a). In parallel, VN-treated cultures exhibited increased total cell numbers compared with controls (Figure 5b). Consistent with enhanced proliferative activity, immunofluorescence analysis revealed a higher proportion of cells positive for the mitotic marker phosphorylated histone H3 (Figure 5c). Notably, VN also promoted cell expansion in three-dimensional suspension cultures, where spheroids derived from chicken embryonic myogenic cells showed increased cellularity in the presence of VN (Figure 5d).

**Figure 5.**
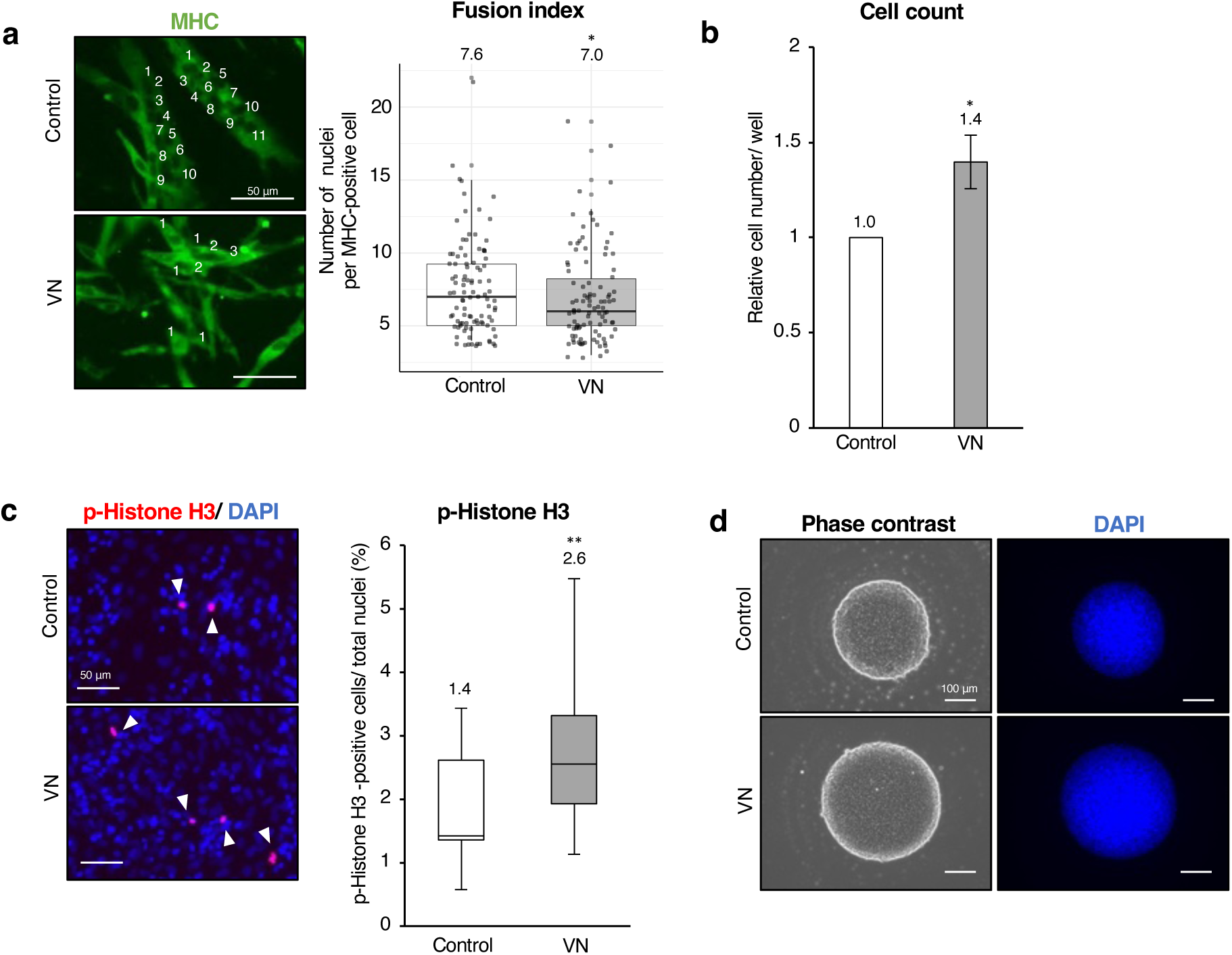
Vitronectin inhibits differentiation and enhances proliferation in primary chicken myoblasts. **a,** Representative immunofluorescence images for MHC showing myotube maturation. Maturation was quantified by the number of nuclei per MHC-positive myotube (Control: n=100, Vitronectin: n = 100 in MHC-positive cells). Scale bar: 50 µm. **b,** The relative cell number of chick myogenic cells cultured in indicated conditions (n = 3). **c,** Representative immunofluorescence images for phosphorylated histone H3 (p-Histone H3) and the proportion of positive cells relative to total nuclei (Control: n = 7619, Vitronectin: n = 7928). Scale bar: 50 µm. **d,** Representative phase-contrast and nuclei images of spheroid on day 5 in rotary suspension spheroid culture. Scale bar: 100 µm. In **a**-**c**, cells were analyzed 6 h after differentiation induction in DM. Nuclei were counterstained with DAPI. Data are presented as mean ± standard deviation. Statistical significance was determined using Student’s t-test; \*\**p < 0.01*.

Together, these results demonstrate that VN-mediated maintenance of a proliferative, differentiation-resistant state is conserved across species and operates not only in immortalized murine myoblasts but also in primary embryonic skeletal muscle cells.

### Vitronectin accumulation links replicative senescence to impaired myogenic differentiation

C2C12 myoblasts progressively lose their differentiation capacity upon long-term passaging, a phenotype associated with replicative senescence (20). Consistent with this, low-passage cells efficiently formed multinucleated, MHC-positive myotubes upon transfer to differentiation medium, whereas differentiation efficiency declined with increasing passage number, and high-passage cultures showed sparse myotube formation (Figure 6a). Immunoblot analysis confirmed robust induction of myogenin and MHC in early-passage cells, which was strongly attenuated in later passages (Figure 6b).

**Figure 6.**
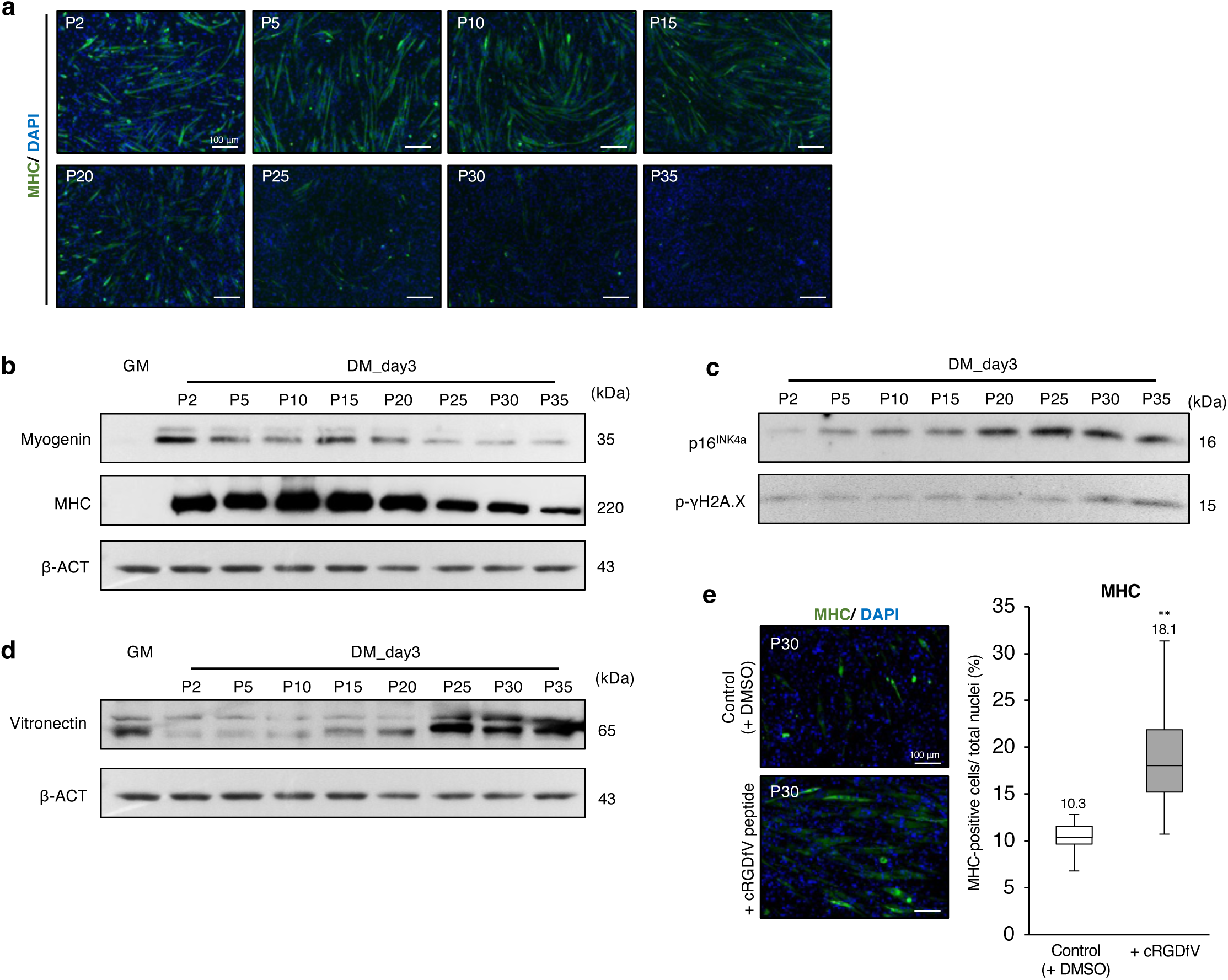
Passage-dependent impairment of myogenic differentiation is accompanied by progressive accumulation of VN in C2C12 cells. **a,** Representative immunofluorescence images showing MHC expression in C2C12 cells cultured in DM for 3 days at indicated passage numbers. Scale bar: 100 µm. **b,** Immunoblot analysis of representative myogenic markers (Myogenin and MHC) in cells across different passages cultured in DM for 3 days. **c,** Immunoblot analysis of senescence-associated markers (p16 and phosphorylated γH2A.X). **d,** Immunoblot analysis for VN protein levels showing progressive accumulation with increasing passage number. **e,** Immunofluorescence analysis of MHC expression in high-passage (P30) C2C12 cells cultured in DM for 3 days under control conditions with DMSO or in the presence of the integrin αvβ3 inhibitor cRGDfV. Quantification shows the percentage of MHC-positive cells relative to total nuclei (differentiation index) (Control: n = 5397, cRGDfV peptide: n = 5105). Scale bar: 100 µm. Nuclei were counterstained with DAPI. Data are presented as mean ± s.d. Statistical significance was determined using Student’s t-test; \*\**p < 0.01*.

Late-passage cells exhibited elevated expression of the senescence markers p16^INK4a^ (49,50) and phosphorylated γH2A.X (51), confirming the establishment of replicative senescence (Figure 6c). Notably, VN protein levels increased progressively with passage number (Figure 6d), despite no corresponding increase in VN mRNA expression (Figure S7), indicating that VN accumulation occurs independently of transcriptional upregulation.

To assess whether VN accumulation functionally contributes to senescence-associated differentiation defects, we inhibited integrin αvβ3, the principal VN receptor, using the competitive peptide cRGDfV in high-passage (P30) cells. Inhibition of αvβ3 signaling partially restored myogenic differentiation, as reflected by increased numbers of MHC-positive cells and a higher differentiation index compared with vehicle controls (Figure 6e).

Together, these results demonstrate that VN accumulates during replicative senescence and functionally contributes to impaired myogenic differentiation through integrin-dependent signaling, identifying an extracellular mechanism linking replicative aging to loss of myogenic competence.

## Discussion

This study identifies vitronectin (VN) as a decisive extracellular regulator that mechanistically links serum composition, growth factor utilization, three-dimensional expansion capacity, and replicative aging to the control of myoblast fate. Our findings demonstrate that VN is not a passive adhesion molecule but an active extracellular signaling platform that sustains proliferative competence while restraining myogenic differentiation, positioning VN as an ECM “gatekeeper” that determines whether myoblasts remain self-renewing or proceed toward terminal commitment.

A central question raised by these findings concerns how VN–integrin αvβ3 signaling suppresses the activation of myogenic differentiation pathways. In our system, VN exposure attenuated phosphorylation of STAT3, ERK, and p38 MAPK pathways (Figure 1e) previously implicated in promoting myogenic differentiation (30,32–34). Whether this inhibition reflects a direct consequence of αvβ3-dependent intracellular signaling or a secondary effect of altered cell cycle dynamics remains unclear. Notably, VN robustly maintained proliferation and prevented cell-cycle exit, a prerequisite for myogenic commitment (Figure 2). Continuous cycling itself is known to antagonize differentiation programs, suggesting that VN-integrin engagement may concurrently transmit anti-differentiation cues and stabilize a proliferative state that indirectly limits full activation of differentiation-associated kinase cascades.

Consistent with this interpretation, VN preserved IGFR and FGFR phosphorylation even under differentiation conditions, suggesting that VN sustains growth factor signaling in environments where it would normally decline (Figure 2f). Integrin signaling has been shown to stabilize membrane-associated signaling platforms and coordinate growth factor receptor clustering (35,52). Previous studies have demonstrated direct and indirect crosstalk between integrins and growth factor receptors, including IGF1R–integrin complexes (53) and integrin-dependent facilitation of FGFR signaling (54). VN may therefore provide an ECM-dependent scaffold that prolongs receptor activation and signaling duration despite reduced availability of soluble mitogens. Elucidating how ECM architecture integrates adhesion receptors with growth factor signaling in myoblasts will be an important avenue for future investigation.

The proliferative and differentiation-suppressive effects of VN were further amplified in three-dimensional rotary spheroid culture, where VN promoted aggregate growth while limiting myotube formation (Figure 4). Premature differentiation represents a major bottleneck in current skeletal muscle organoid and 3D culture systems, restricting progenitor expansion and tissue maturation (55,56). Our findings suggest that VN-rich extracellular environments may stabilize a proliferative amplification phase prior to differentiation. Importantly, comparable effects were observed in primary embryonic chicken myogenic cells (Figure 5), indicating that VN-mediated gatekeeping is not restricted to immortalized murine cells but instead reflects a conserved regulatory principle across species.

Our serum-free culture experiments further demonstrate that VN alone was insufficient to sustain long-term myoblast survival and expansion; instead, cooperative signaling inputs were required (Figure 4g; Figure S4). Among these, LIF emerged as an essential component, as VN-supported serum-free cultures failed to maintain viable, proliferative cells in its absence. Although LIF was initially introduced on the basis of serum-free mouse embryonic stem (ES) cell media formulations (26), its essential requirement is consistent with previous observations that LIF can modulate myogenic cell fate in a context-dependent manner (29), supporting survival and proliferation while restraining premature differentiation (57,58). Importantly, LIF is not considered a canonical mitogen for myoblast proliferation, distinguishing its role from classical growth factors such as FGF2 or IGF-1.

In contrast, cholesterol and 2-mercaptoethanol (2-ME) functioned primarily as supportive factors that enhanced proliferative robustness rather than determining survival. Cholesterol likely stabilizes membrane architecture and signaling microdomains that facilitate growth factor receptor organization and signaling efficiency (59), whereas 2-ME may improve cellular redox balance and stress tolerance (60). Exogenous FGF2 could further augment proliferation, indicating that VN provides a permissive adhesive framework upon which additional pro-mitogenic cues can be layered. Importantly, cells expanded under VN–LIF–based serum-free conditions retained full differentiation competence when subsequently exposed to HS-containing differentiation medium (Figure S5), demonstrating that this culture system preserves a reversible, proliferation-competent state rather than driving irreversible dedifferentiation or exhaustion.

Another major implication of our findings concerns replicative senescence (Figure 6). VN progressively accumulated during extended passaging, in parallel with declining differentiation competence and increasing senescence markers such as p16 and phosphorylated γH2A.X. These observations suggest that long-term VN exposure contributes to the erosion of myogenic capacity characteristic of replicative aging. The mechanism underlying VN accumulation remains unsolved; however, it is unlikely to be explained by transcriptional upregulation, because VN protein elevation occurs without a corresponding increase in *VN* gene expression (Figure S7). These findings raise the possibility that non-transcriptional mechanisms, such as reduced turnover, impaired matrix remodeling, or altered feedback regulation of secreted VN, may account for its progressive accumulation with passage. Interestingly, although VN supports proliferation, senescent cultures ultimately lose proliferative capacity, reflecting the emergence of intrinsic aging constraints such as p16-mediated cell-cycle arrest and DNA damage accumulation that override VN’s pro-mitogenic influence (61–63). Importantly, blocking αvβ3 signaling partially restored differentiation in high-passage cells, directly implicating VN–integrin signaling in senescence-associated differentiation failure. Given that late-passage C2C12 cells have been shown to recapitulate key features of aged skeletal muscle (64), these findings raise the possibility that VN accumulation contributes to age-related declines in regenerative capacity. Future studies should therefore examine whether VN similarly accumulates in aged or sarcopenic muscle tissue and whether targeting VN–integrin signaling can restore myogenic plasticity in aging contexts.

In summary, our study establishes VN as an ECM gatekeeper that integrates adhesion signaling, growth factor receptor activity, serum composition, three-dimensional culture behavior, and replicative aging into a unified framework governing myoblast fate. These findings refine long-standing assumptions about serum-dependent myogenesis and highlight extracellular matrix composition as a critical, and previously underappreciated, determinant of muscle cell fate decisions.

## Acknowledgments

We would like to thank Prof. Yoko Kato (Kindai Univ.) for the valuable comments and Jafar Sharif (Laboratory for Developmental Genetics, RIKEN Center for Integrative Medical Sciences (IMS)) for the kindly gifted materials.

